# Mitral valve leaflet response to ischemic mitral regurgitation: From gene expression to tissue remodeling

**DOI:** 10.1101/864876

**Authors:** Daniel P. Howsmon, Bruno V. Rego, Estibaliz Castillero, Salma Ayoub, Amir H. Khalighi, Robert C. Gorman, Joseph H. Gorman, Giovanni Ferrari, Michael S. Sacks

## Abstract

**Aims:** Ischemic mitral regurgitation is frequently observed following myocardial infarction and is associated with higher mortality and poor clinical prognosis if left untreated. Accumulating evidence suggests that mitral valve leaflets actively remodel post–myocardial infarction, yet the cellular mechanisms underlying these responses and how this affects tissue function remain largely unknown. We sought to elucidate mitral valve remodeling post myocardial infarction at the tissue, cellular, and transcriptomic levels.

**Methods and Results:** The mechanical behavior of ovine mitral valve leaflets pre– and 8 weeks post– myocardial infarction reveal a significant decrease in radial direction extensibility, which essentially eliminated the mechanical anisotropy typically observed in healthy mitral valves. Quantitative histology and ultrastructural assessment by transmission electron microscopy revealed altered leaflet composition and architecture at 8 weeks post–myocardial infarction. Assessment of the mitral valve interstitial cell nuclear aspect ratio, a metric of cellular deformation, revealed that they were on average rounder following myocardial infarction. RNA sequencing indicated that YAP-induced genes were elevated at 4 weeks post–myocardial infarction and genes related to extracellular matrix organization were some of the most downregulated in sheep with IMR compared to sheep without ischemic mitral regurgitation at 4 weeks post–myocardial infarction. Additionally, RNA sequencing revealed the possible recruitment of immune cells in this remodeling process due to the drastic elevation of CXCL9 and CLEC10A.

**Conclusions:** Our multiscale assessment revealed significant mechanical and microstructural changes due to myocardial infarction. RNA sequencing provided a baseline for global gene expression changes in response to myocardial infarction with and without ischemic mitral regurgitation and suggests YAP-induced mechanotransduction, altered expression of extracellular matrix–related genes, and recruitment of immune cells as mechanisms contributing to altered mitral valve biomechanics post–myocardial infarction.

## Introduction

The fundamental physiological function of the mitral valve (MV) is to facilitate unidirectional blood flow from the left atrium to the left ventricle (LV). Both the MV anterior and posterior leaflets (MVAL and MVPL, respectively) are connected to the papillary muscles (PMs) via the chordae tendineae (MVCT), creating an intimate mechanical coupling with the LV.^1–3^ MV pathologies are most commonly associated with inadequate leaflet coaptation during systole, which leads to mitral regurgitation (MR) and subsequent retrograde flow into the left atrium. MR leads in turn to elevated risk of pulmonary congestion, heart failure, and stroke. Moreover, MR is the most common valvular heart disease in developed countries, with an estimated prevalence of 1.7%.^4^ Whereas primary MR results from an intrinsically diseased valve (e.g., myxomatous disease), secondary MR occurs in response to adverse remodeling of the LV (e.g., due to ischemic cardiomyopathy) due to the intimate coupling between the MV and LV. Ischemic MR (IMR), secondary MR due to ischemic cardiomyopathy, is present after myocardial infarction (MI) in 40–50% of cases^5,6^ (with moderate/severe IMR in 12–19% of cases^5,7^). LV remodeling begets IMR which begets further LV remodeling^8^, creating a positive feedback loop that doubles the rate of cardiac mortality post-MI when IMR is present.^7,9,10^

IMR is treated surgically either through MV repair or replacement, with repair currently the preferred method under the assumption that preserving the normal cardiac LV-MV structural interactions is beneficial. Early reports regarding MV replacement revealed a higher 30-day postoperative risk of death compared to valve repair (OR = 2.52, 95%CI 1.91–3.34) for severe IMR.^11^ Moreover, repair without recurrent IMR results in superior LV volume reduction over replacement, but this LV volume reduction vanished if IMR recurred.^12^ However, more recent studies in patients with severe IMR report similar death rates for repair vs. chordal-sparing replacement, but the IMR recurrence rate was much higher in the repair group (32.6% vs. 2.3%) at 1 year^12^ and (58.8% vs. 3.8%) at 2 years^13^ post-surgery. Newer, less invasive surgical techniques, such as the MitraClip, promise to address both the safety of the procedure and the IMR recurrence rate^14^; however, long-term outcomes for these approaches remain unknown. In particular, the state of the MV leaflet tissue at the time of intervention remains a major unknown. Given the fact that interventions such as the MitraClip induce stress concentrations at the attachment points, it is likely that remodeling in these areas may be of greater magnitude than those induced either by IMR or annuloplasty techniques.

Although IMR is classically depicted as functional, (i.e., without intrinsic changes to the MV), accumulating evidence suggests that the MV undergoes substantial remodeling post-MI. This is not surprising since it has been demonstrated that nonpathologic LV volume overload induced by pregnancy results in leaflet tissue remodeling, including increases in leaflet size and relative collagen content.^15–18^ Total MV area, MV thickness, and collagen content increased in both the human MV^19^ and in ovine models^20,21^ of IMR. Further, in an ovine model with tethered leaflets to emulate the effects of IMR without the confounding effects of LV ischemia, the MV exhibited increases in area and thickness^22^, indicating that leaflet tethering is likely driving this remodeling. MV remodeling appears to be regionally heterogeneous; our group has recently demonstrated in an ovine model that post-MI the MV leaflets are permanently distended in the radial direction, with the magnitude and rate of distention varying regionally over the leaflet surface.^23^

Regardless of the exact causes, continued MV remodeling in MR (especially the onset and progression of fibrosis and calcification) is thought to underlie reported repair failures.^24^ These findings also agree with computed tomography assessments that observe thickened leaflets and mitral stenosis in a subset of repaired MVs.^25^ Clearly, a better understanding of MV remodeling post-MI directly ties into the clinical success of current surgical interventions. As in all heart valves, the valve interstitial cells (VICs) are the primary drivers of valve tissue growth and remodeling in response to both biochemical and mechanical stimuli.^26^ MV tissue culture in a circumferentially-oriented strip bioreactor after 48 hours of cyclic strain significantly upregulated many genes related to ECM remodeling.^27^ Altered mechanical loads following undersized ring annuloplasty and/or papillary muscle approximation also show differences in several genes related to ECM remodeling.^28^ This feedback interaction between VICs and their micro-mechanical environment and the way this scales to the levels of the tissue and organ (valve and ventricle) warrants further investigation, especially in the context of IMR.

Although recent reports have continued to shed light on the MV remodeling processes post-MI, either unaltered^23,29,30^ or in the context of various interventions^25,28,31^, fundamental questions regarding MV remodeling in IMR remain. In particular, detailed analyses of the functional physiological behaviors of the post-MI MV necessary for informing advanced computational simulations of patient-specific surgical procedures remain lacking. At the underlying sub-cellular response level, individual genes/proteins (e.g.,^29,31^) or gene panels related to a single process^28^ have been investigated, but there has not been a transcriptomics study that investigates all genes/processes that change in response to IMR in vivo. Transcriptomics provides an unbiased approach for investigating the importance of individual genes/processes in the context of global gene expression, potentially uncovering novel genes that would otherwise be overlooked by deciding which genes should be investigated a priori.

The present study thus sought to address these shortcomings through a detailed investigation of the impact of IMR at the transcriptomic, cellular, and tissue levels in an established ovine model of IMR. MV leaflet tissue explants were evaluated to assess the effects of IMR, including changes in the leaflet tissue mechanical response, quantitative histology to investigate the extracellular matrix (ECM), and VIC nuclear aspect ratio (NAR) to quantify cell-level geometric changes, which has been associated with VIC biosynthetic levels. In addition, RNA sequencing was used to quantify global gene expression changes post-MI in IMR and normal groups. By connecting functional behavior of the MV leaflets at the highest scale down to changes in gene expression at the lowest scale, this study provides crucial knowledge for designing next-generation surgical devices and suggesting possible points of pharmaceutical intervention to improve the current dismal clinical outcomes for treating IMR.

## Methods

### 1. IMR ovine model

All animal protocols used in this study were approved by the University of Pennsylvania’s Institutional Animal Care and Use Committee and complied with the National Institute of Health’s guidelines for the care and use of laboratory animals (NIH Publication 85-23, revised 1996). The University of Pennsylvania, with the supervision of the School of Veterinary Medicine, maintains a full-service vivarium facility operated by the University Laboratory Animal Research (ULAR) organization. The vivarium is directly adjacent to the laboratory’s operating room suite.

IMR was induced in adult (30–40 kg) Dorset sheep raised for laboratory work and supplied by commercial vendors following established procedures.^32–34^ Briefly, sheep were anaesthetized with sodium thiopental (10–15 mg/kg) intravenously, intubated, and ventilated with isoflurane (1.5–2%) and oxygen. Surface electrocardiogram, arterial blood pressure, and other vital signs were continuously monitored throughout the duration of the procedure. IMR was induced by a thoracotomy to facilitate ligation of the second and third obtuse marginal branches of the circumflex coronary artery. Permanent occlusion of these arteries reliably results in a transmural posterior MI that includes the entire posterior PM and involves approximately 20% of the LV mass, causing a gradual onset of severe IMR within eight weeks. The degree of MR was assessed with echocardiography. MV leaflets were collected and flash-frozen at pre-MI as well as 4 and 8 weeks post-MI. All sheep were euthanized by an overdose of KCl and sodium thiopental while under general anesthesia. The animals were pronounced dead only after ECG silence is demonstrated for 3–5 minutes and cardiac arrest is visibly confirmed. This technique has been approved by the University of Pennsylvania IACUC and is consistent with the recommendations of the American Veterinary Medical Association (AVMA) Guidelines on Euthanasia. Leaflets from one subset of 14 animals were used for mechanical testing (pre-MI, n=6; 8 weeks post-MI, n=8) and tissue characterization via histology and electron microscopy (pre-MI, n=3; 8 weeks post-MI, n=8). All of the animals in this first subset displayed low-grade IMR at 8 weeks post-MI. Leaflets from a second subset of 13 animals (pre-MI, n=4; 4 weeks post-MI, n=5; 8 weeks post-MI, n=4) were used for RNA isolation and quantification. Half of the animals at 4 and 8 weeks post-MI exhibited low-grade IMR, with the other half exhibiting no observable IMR. No animals exhibited IMR at the pre-MI time point.

### 2. MV leaflet tissue mechanical evaluation

As the MV leaflets are relatively thin, their mechanical behavior has been extensively characterized using planar biaxial mechanical testing.^35–37^ In the present study, we utilized our established experimental biaxial system to evaluate the mechanical behaviors of 1 × 1 cm^2^ specimens extracted from the upper mid regions of 14 ovine MV anterior leaflets (pre-MI, n=6; 8 weeks post-MI, n=8) following established procedures.^36,37^ Specifically, each specimen was loaded to an approximate end-systolic peak leaflet tissue stress of 200 kPa, while setting the following circumferential-to-radial ratios of the first Piola–Kirchhoff stress tensor components *P*_CC_: *P*_RR_ to values of 1:2, 3:4, 1:1, 4:3, and 2:1. For each specimen, the right Cauchy–Green deformation tensor ***C*** and second Piola–Kirchhoff stress tensors ***S*** were computed at each instant of loading. For clarity, an illustration of the location of the samples for mechanical testing and ultrastructural analysis are provided in Figure 1.

**Figure 1:**
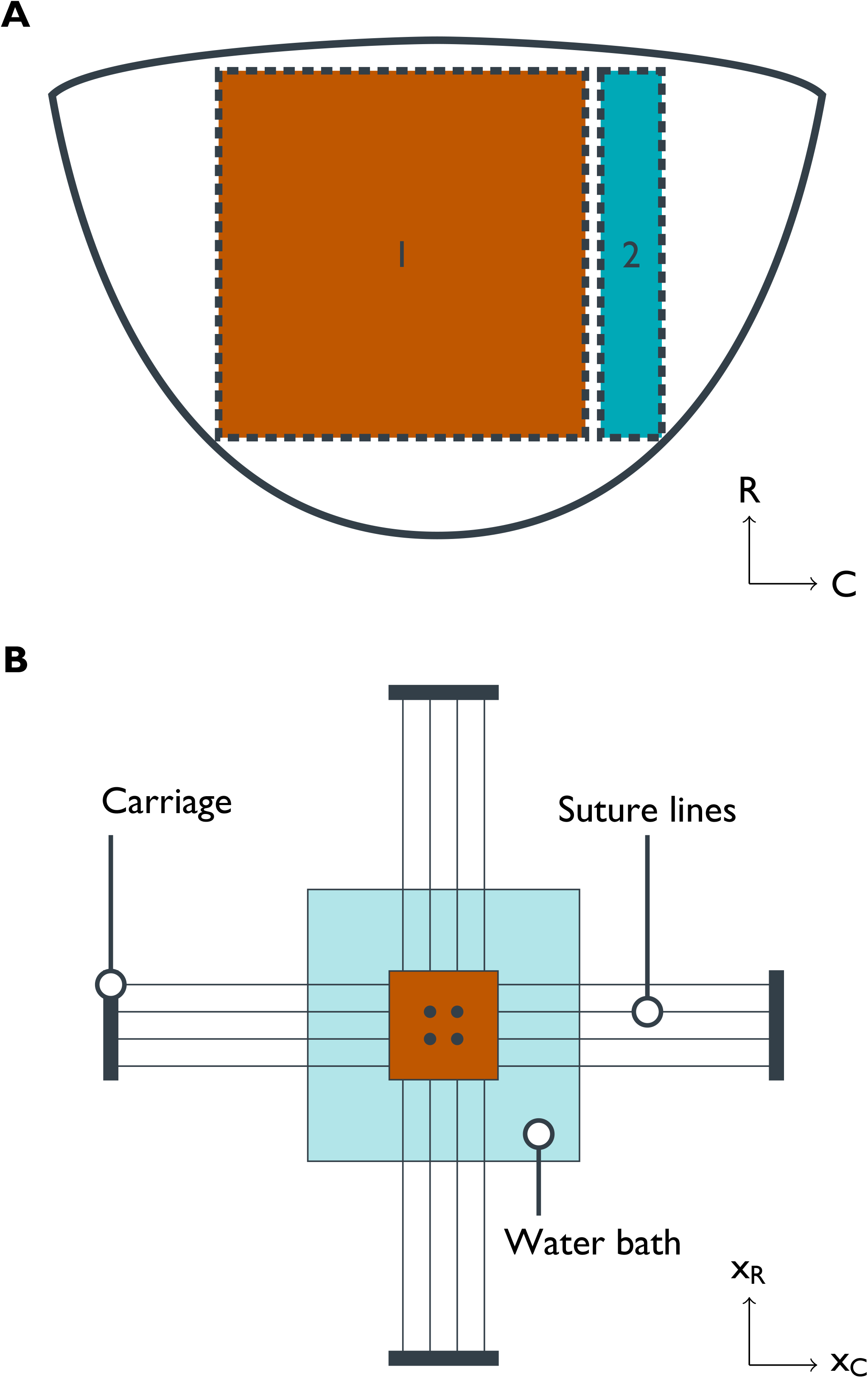
Tissue preparation for mechanical testing and tissue-level experiments. (A) Illustration of tissue explant with radial (R) and circumferential (C) directions used for mechanical testing (region 1) and that used for histology and TEM (region 2). (B) Illustration of the biaxial mechanical testing apparatus, with four centrally located fiducial markers, sutures mounting the tissue sample, and displacements *x_R_* and *x_c_* in the radial and circumferential directions, respectively.

One of the major goals of the present study was to reveal changes in intrinsic mechanical behavior of the primary structural ECM constituent collagen fibers. However, this cannot be derived directly from the mechanical data, since alterations in tissue-level behavior can be a combined result of changes in collagen stiffness and in their physical arrangement. To separate these two effects, we thus utilized a structural constitutive (material) model, as presented and applied extensively for heart valve tissues.^18,38,39^ Briefly, we applied the following modeling approach to the measured stress-strain data using

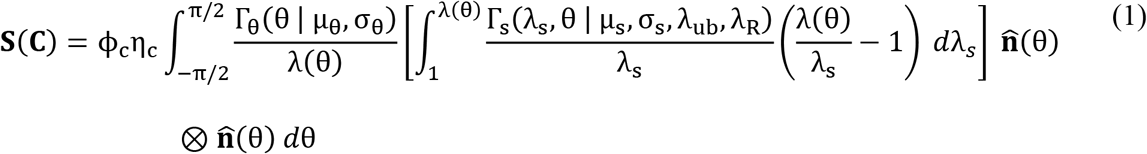

where ϕ_c_ is the collagen mass fraction (measured via histology); η_c_ is the estimated collagen fiber modulus; Γ is the collagen orientation distribution as a function of the in-plane angle θ, with mean μ and standard deviation σ; 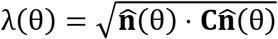 is the stretch along the direction vector 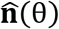 and Γ_s_ is the angle-dependent distribution of the collagen fiber slack stretch λ_s_, with mean μ_s_, standard deviation σ_s_, and upper bound λ_ub_ in the circumferential direction (i.e. at θ = 0).

Although in normal MV leaflet tissues, the collagen fiber slack stretch λ_s_ is largely independent of angle^39^, MI has been shown to induce large leaflet distentions along the radial direction^23^. Such changes are sufficient to make the distribution Γ_s_ dependent on θ, even in the absence of active cell-mediated remodeling. To account for this effect and distinguish it from structural changes due to active remodeling, we introduced the radial pre-stretch parameter λ_R_, which quantifies the radial stretch necessary to make Γ_s_ independent of angle. At any angle θ, the corresponding pre-stretch is 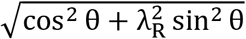. Therefore, with respect to the reference configuration for mechanical testing, the angle-specific mean, standard deviation, and upper bound of Γ_s_ are μ_s_/λ, σ_s_/λ, and λ_ub_/λ respectively.

A key feature of this formulation is that the model parameters have a *direct physical meaning.* In particular, we note that changes in collagen fiber modulus η_c_ and the two structural functions Γ and Γ_s_ can be separated. This allows us to address the question of whether changes due to IMR alter the intrinsic collagen fiber modulus (due to fatigue damage or active cell-mediated remodeling), the fibrous structure, or both.

## Histology

### 3. Histological preparation

Tissue preparation for histology has been described previously.^40,41^ Briefly, samples from MV anterior leaflets (both pre-MI and 8 weeks post-MI) were fixed with 10% buffered formalin, embedded in paraffin, cut into 5 μm sections, and mounted on glass slides. Slides were then stained with either Movat’s Pentachrome for visualization of collagen (yellow-orange), glycosaminoglycans (GAGs; light blue), and elastin (dark purple) or hematoxylin and eosin (H&E) for determination of cell density and nuclear geometry. Slides were imaged using a light microscope at 4×, 10×, and 20× magnification (Leica, Wetzlar, Germany).

### 4. Quantitative analysis

The relative presence and distribution of each ECM constituent was quantified directly from the resulting images using a pixel-wise color analysis protocol, extended from methods presented previously.^42^ First, the Movat image was converted from a red-green-blue (RGB) format to a hue-saturation-value (HSV) format. Then, the hue distribution of the tissue-occupied pixels was fit to a three-component wrapped Gaussian mixture model via maximum likelihood estimation.

To describe the distribution of collagen, GAG, and elastin mass fractions through the thickness of the leaflet, the tissue-occupied region of each Movat image was transformed to a rectangular domain using a conformal (locally angle-preserving) map such that coordinate directions in the new domain represent the circumferential and transmural directions of the leaflet (Figure 2), similar to a previous method for thickness characterization.^43^ This was accomplished through the following procedure: First, the top and bottom boundaries of the tissue region were determined automatically using the saturation mask previously defined to isolate the tissue-occupied region and traced from end to end. Second, local circumferential direction vectors on the boundaries were defined as the unit tangent to the boundary at each point. Third, the circumferential direction vector components were interpolated using a 2D spline function to define the circumferential vector field everywhere within the tissue region. Finally, transmural direction vectors were determined through a 90-degree rotation of the local circumferential direction. Streamlines of the transmural vector field were then traced from boundary to boundary and discretized with segments of equal arc length, in order to express the thickness-dependent variation in tissue composition with respect to a normalized transmural coordinate (NTC). Transmural tracing was performed sequentially at circumferential locations that were equally spaced halfway through the tissue thickness (defined by the midpoints of the traced transmural streamlines, in an arc length sense).

**Figure 2:**
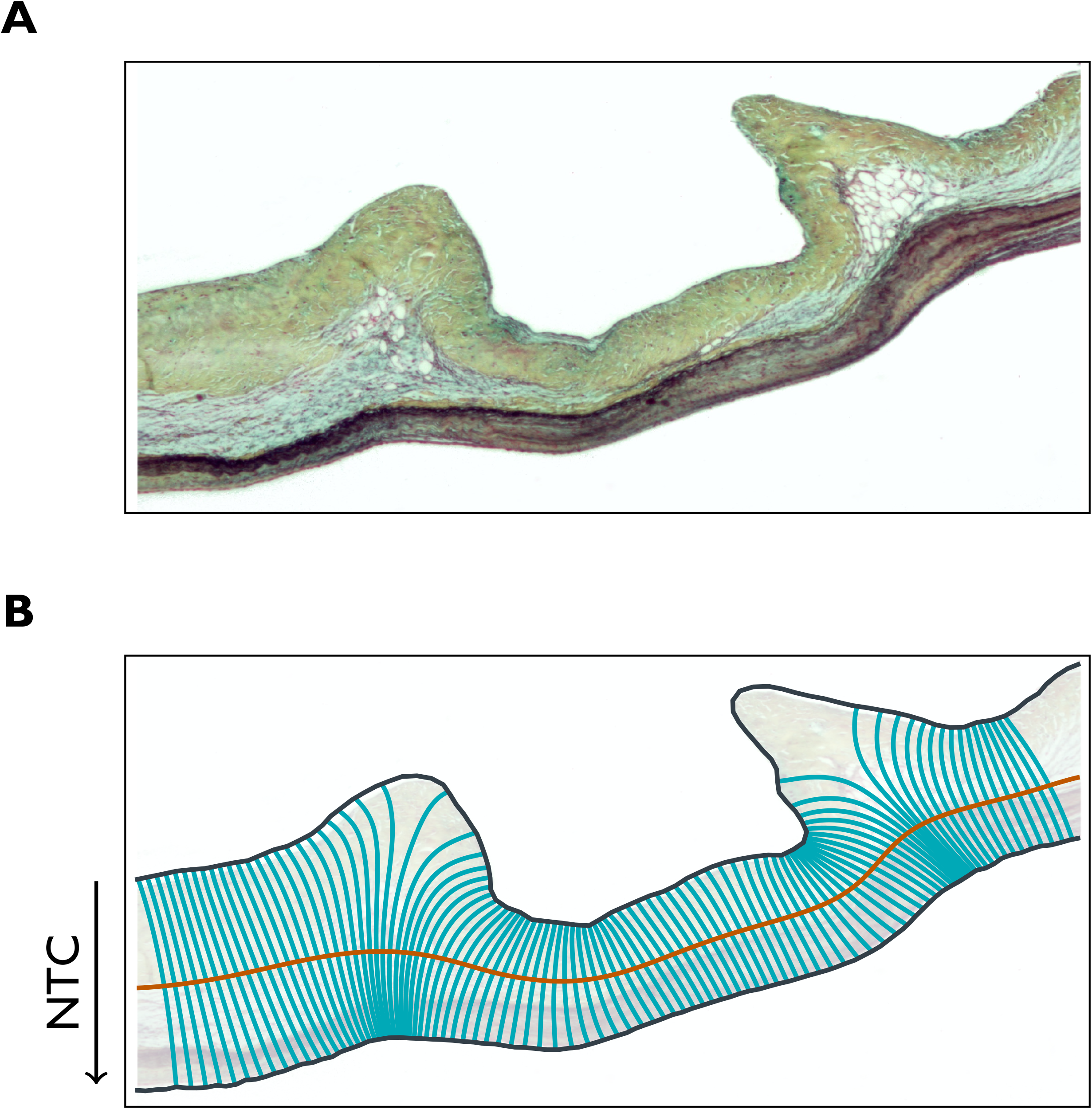
Example of (A) Movat stained sample and (B) the corresponding conformal coordinate system with the direction of the NTC.

Since the ECM compositions (in 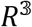) are constrained such that they sum to unity, the mass fractions were transformed to an orthonormal coordinate system in 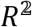 using compositional data analysis.^44^ Anderson-Darling tests for normality and Hotelling’s T^2^ test for differences in location between two distributions were then carried out in the orthonormal coordinate space.

### 5. Determination of the nuclear aspect ratio (NAR)

H&E-stained slides, which provide a sharp contrast between nuclei and the surrounding collagen fibers, were used to determine MVIC NAR. A custom-written MATLAB (MathWorks, Inc., Natick, MA) script was used to automatically detect and fit ellipses to 10 nuclei in each H\&E image and the NAR of each MVIC was then calculated as the ratio between the major and minor radii of the corresponding ellipse.

### 6. Ultrastructural characterization via transmission electron microscopy (TEM)

To evaluate changes in the architecture of the ECM owing to MI, samples were imaged using transmission electron microscopy (TEM). Samples were fixed in Grade II EM Grade Glutaraldehyde (Electron Microscopy Sciences, Hatfield, PA) for a total of four hours, then stained with osmium tetroxide for four hours, dehydrated with alcohol and acetone, infiltrated in epoxy resin overnight, and cured for 48 h in a 60°C oven. The resulting resin blocks were trimmed with a razor blade into a trapezoid block face such that the y-axis represents the transmural/thickness axis and the x-axis represents the circumferential axis of the MVAL. Thin sections (70 nm) were cut using a Diatome Ultra Diamond Knife 35° (Diatome Diamond Knives, Hatfield, PA), picked up with Formvar-coated slot grids (Electron Microscopy Sciences, Hatfield, PA), and imaged with a FEI Tecnai TEM (FEI, Hillsboro, OR).

### 7. RNA sequencing (RNA-seq) workflow

#### 7.1 RNA isolation

Isolation of total mRNA was performed using the RNeasy mini kit (Qiagen, Valencia, CA). 1% β-mercaptoethanol was added to the lysis buffer and 30 mg of previously flash-frozen frozen tissue were homogenized with a TissueRuptor (Qiagen). Isolated RNA purity was preliminarily assessed on a NanoDrop spectrophotometer (NanoDrop technologies, Wilmington, DE). RNA concentration was measured on a Qubit 2.0 fluorometer (Invitrogen, Whaltham, MA). RNA integrity assessment was performed on an Agilent Bioanalyzer 2100 using a Nano 6000 assay kit (Agilent Technologies, Santa Clara, CA). A RNA integrity number (RIN) > 7.1 was considered the minimum requirement for library preparation. RNA from one anterior leaflet did not meet this requirement and was removed from analysis. All remaining samples displayed RINs > 8.6.

#### 7.2 RNA library construction and transcriptome sequencing

RNA library preparation and sequencing were performed at Novogene (Sacramento, CA). RNA quality control by 1% agarose gels and Agilent Bioanalizer 2100 was repeated before library construction. A total amount of 1 μg RNA per sample was used as input material for the RNA sample preparations. Sequencing libraries were generated using NEBNext^®^ Ultra™ RNA Library Prep Kit for Illumina^®^ (NEB, USA) following manufacturer’s recommendations and index codes were added to attribute sequences to each sample. Briefly, mRNA was purified from total RNA using poly-T oligo-attached magnetic beads. Fragmentation was carried out using divalent cations under elevated temperature in NEBNext First Strand Synthesis Reaction Buffer (5×). First strand cDNA was synthesized using random hexamer primer and M-MuLV Reverse Transcriptase (RNase H-). Second strand cDNA synthesis was subsequently performed using DNA Polymerase I and RNase H. Remaining overhangs were converted into blunt ends via exonuclease/polymerase activities. After adenylation of 3’ ends of DNA fragments, NEBNext Adaptor with hairpin loop structure were ligated to prepare for hybridization. In order to select cDNA fragments of preferentially 150–200 bp in length, the library fragments were purified with AMPure XP system (Beckman Coulter, Beverly, USA). 3 μL USER Enzyme (NEB, USA) was used with size-selected, adaptor-ligated cDNA at 37°C for 15 min followed by 5 min at 95°C before PCR. PCR was performed with Phusion High-Fidelity DNA polymerase, Universal PCR primers, and Index (X) Primer. At last, PCR products were purified (AMPure XP system) and library quality was assessed on the Agilent Bioanalyzer 2100 system. The clustering of the index-coded samples was performed on a cBot Cluster Generation System using PE Cluster Kit cBot-HS (Illumina) according to the manufacturer’s instructions. After cluster generation, the library preparations were sequenced on an Illumina platform and 125 bp/150 bp paired-end reads were generated.

#### 7.3 Data preprocessing

Raw reads (in FASTQ format) were processed with Perl scripts. Low quality reads and those containing adapters or poly-N sequences were removed from raw data. The *Ovis aries* v3.1 reference genome was obtained from Ensembl (release 76). The index of the reference genome was built using Bowtie (v2.2.3) and paired-end reads were aligned to the reference genome using TopHat (v2.0.12). HTSeq (v0.6.1) was used to count the reads numbers mapped to each gene.

#### 7.4 Annotation, statistical modeling, and gene set enrichment analysis

All annotation and analysis was performed with BioConductor.^45^ Genes were annotated with NCBI Entrez IDs, gene symbols, human homolog Entrez IDs, and human homolog gene symbols using biomaRt.^46^ Statistical modeling was performed with DESeq2^47^ with shrinkage of log_2_ fold changes provided by the “apeglm”^48^ method. Differential expression (single model) was assessed with a Wald test whereas the comparison of two (nested) models was assessed with a likelihood ratio test (LRT). Multiple hypothesis correction was performed by controlling the false discovery rate (FDR) via Benjamini-Hochberg.^49^ Gene set enrichment analysis^50^ was performed with the Reactome^51^ database via ReactomePA^52^, where genes were described by human homolog Entrez IDs because Reactome does not currently support analysis for sheep. Genes were ranked by their log_2_ fold change after shrinkage. Multiple hypothesis correction was again performed by controlling the false discovery rate (FDR) via Benjamini-Hochberg.^49^

#### 7.5 Data availability

All data for the RNA-seq experiments presented in this publication have been deposited in NCBI’s Gene Expression Omnibus^53^, in compliance with the Minimum Information about a high-throughput nucleotide SEQuencing Experiment (MINSEQE) guidelines, and are accessible through GEO Series accession number GSE139921 (https://www.ncbi.nlm.nih.gov/geo/query/acc.cgi?acc=GSE139921).

## Results

### 1. Mechanical responses

Consistent with previous studies^35–37,54^, leaflets from pre-MI MVs exhibited greater extensibility in the radial direction compared to the circumferential direction (radial peak strain: 18.7% ± 4.7%; circumferential peak strain: 2.9% ± 0.8%). Interestingly, substantial radial stiffening eliminated this anisotropy in the 8-week post-MI group, on average (radial peak strain: 4.9% ± 1.3%; circumferential peak strain: 5.1% ± 0.6%) (Figure 3 and Supplemental Figure 1).

**Figure 3:**
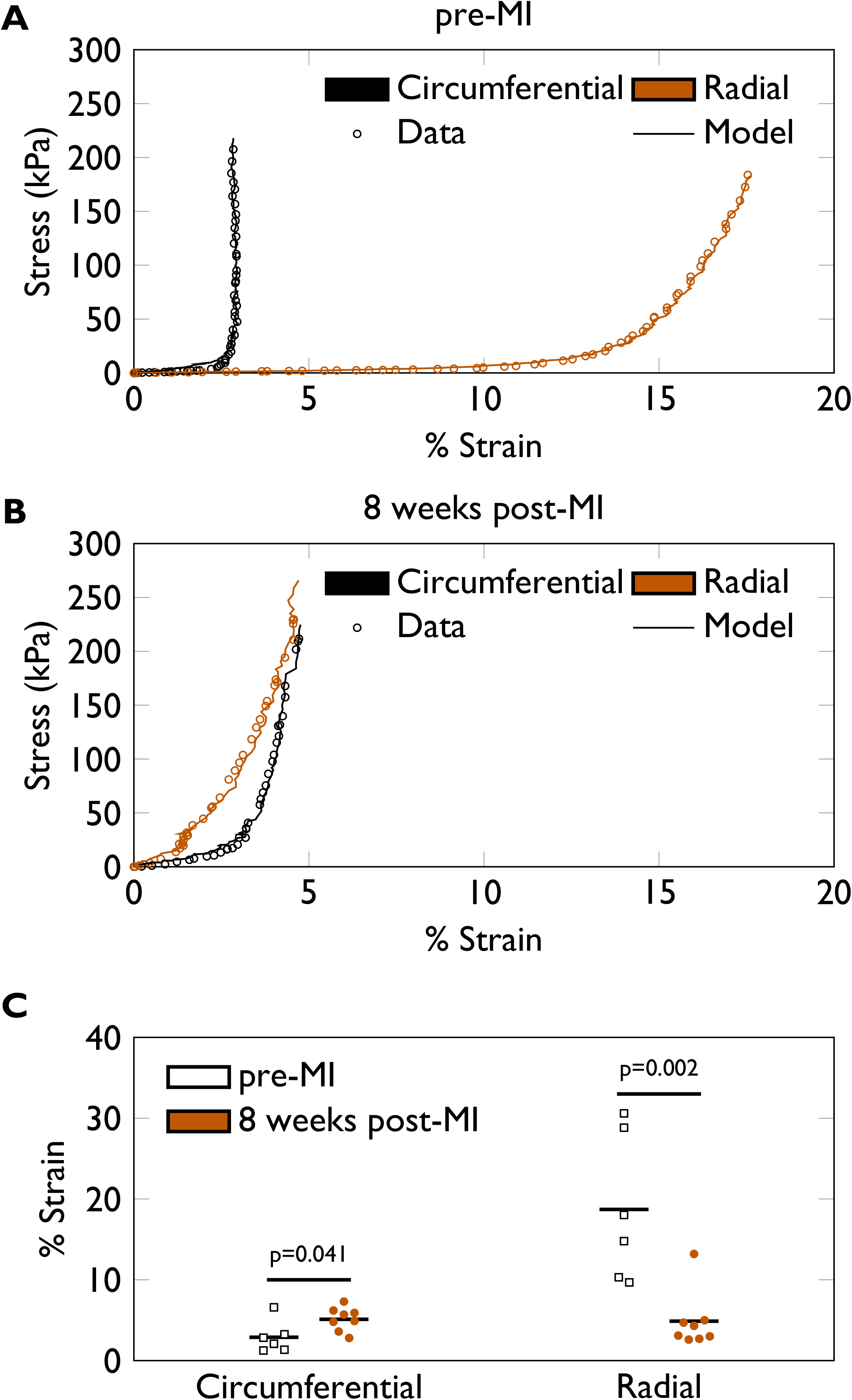
Biaxial mechanical testinga results indicate a loss of mechanical anisotropy at 8 weeks post-MI. Representative stress/strain curves for (A) pre-MI and (B) 8 weeks post-MI specimens, with every third data point plotted to facilitate visualization. (C) Strains at 200 kPa for all specimens tested (pre-MI, n=6; 8 weeks post-MI, n=8). Groups were compared with a Student’s t-test.

Parameter estimates obtained from the structural constitutive model suggest that the observed changes in mechanical properties can be attributed predominantly to MI-induced *permanent* radial distention. Specifically, a statistically significant difference between pre-MI and 8 weeks post-MI groups was only detected in the λ_R_ parameter, which estimates post-MI radial distention (p-value = 0.0003). Our results suggest that collagen fiber crimp is largely independent of angle in pre-MI leaflets (i.e. λ_R_ ≈ 1), consistent with previous findings.^39^ Notably, the estimated mean increase in λ_R_ from pre-MI to 8 weeks post-MI (7–9%; Table 1) agreed well with previous echocardiography-based measurements of radial leaflet lengthening post-MI^23^, where the central region of the anterior leaflet was reported to distend about 10% after 8 weeks. No differences were found with regard to collagen fiber stiffness (η_c_), collagen fiber crimp (μ_s_, σ_s_, λ_ub_), or collagen fiber orientation (μ, σ), beyond those which were induced by passive distention. This suggests that while the fiber architecture was substantially altered due to MI, the collagen fibers themselves are essentially unchanged.

**Table 1:** Summary of parameter estimates from mechanical data. Groups compared with Student’s t-tests.

| Parameter | pre-MI | 8 weeks post-MI | p-value |
| --- | --- | --- | --- |
| $\eta_c$ (MPa) | $213 \pm 27$ | $194 \pm 13$ | 0.50 |
| $\mu_s$ | $1.10 \pm 0.01$ | $1.09 \pm 0.01$ | 0.53 |
| $\sigma_s$ | $0.025 \pm 0.006$ | $0.021 \pm 0.003$ | 0.50 |
| $\lambda_{ub}$ | $1.16 \pm 0.02$ | $1.14 \pm 0.01$ | 0.41 |
| $\lambda_R$ | $0.98 \pm 0.01$ | $1.07 \pm 0.01$ | 0.0003 |
| $\mu$ (deg) | $-6.3 \pm 10.7$ | $1.9 \pm 5.6$ | 0.47 |
| $\sigma$ (deg) | $19.5 \pm 5.5$ | $23.6 \pm 1.4$ | 0.43 |

### 2. Altered fiber architecture and ECM composition at 8 weeks post-MI

Transmission electron micrographs of MV anterior leaflets show visible damage to the collagen fibrillar network following infarction (Figure 4). At 8 weeks post-MI, the ECM exhibited qualitatively shorter collagen fibrils that were largely unbound, possibly as a result of overload-induced collagen rupture. Quantitative histology indicates significant alterations in the overall composition (p-value = 0.001), with generally decreased GAG and increased collagen mass fractions at 8 weeks post-MI (Figure 5). Thus, both the fiber architecture and ECM composition are altered at 8 weeks post-MI.

**Figure 4:**
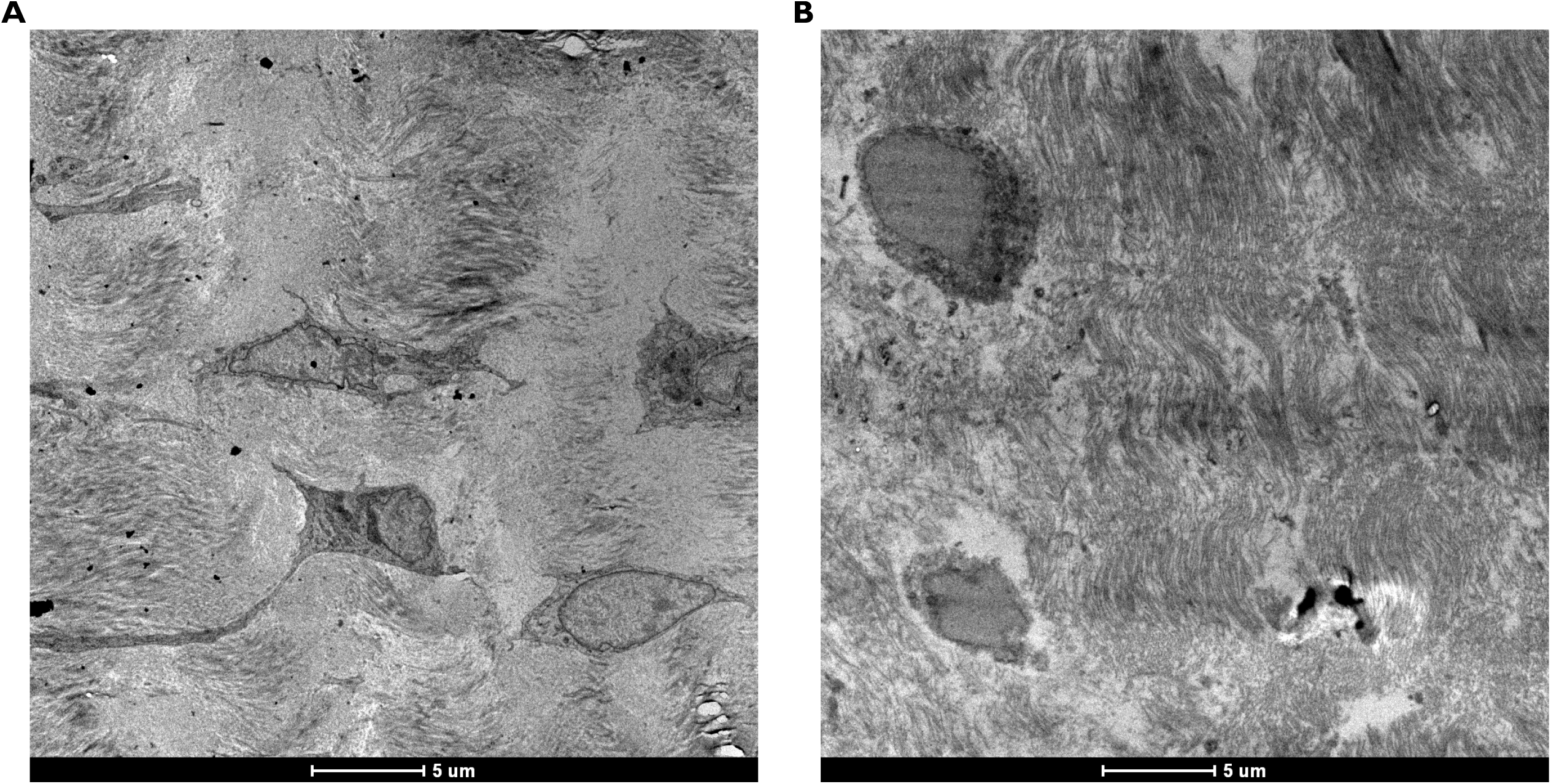
Tissue micro-morphology. Representative micrographs for sample (A) pre-MI and (B) 8 weeks post-MI. Post-MI ECM exhibits a collection of shorter-than-normal fibrils that are largely unbound and VICs appear more rounded.

**Figure 5:**
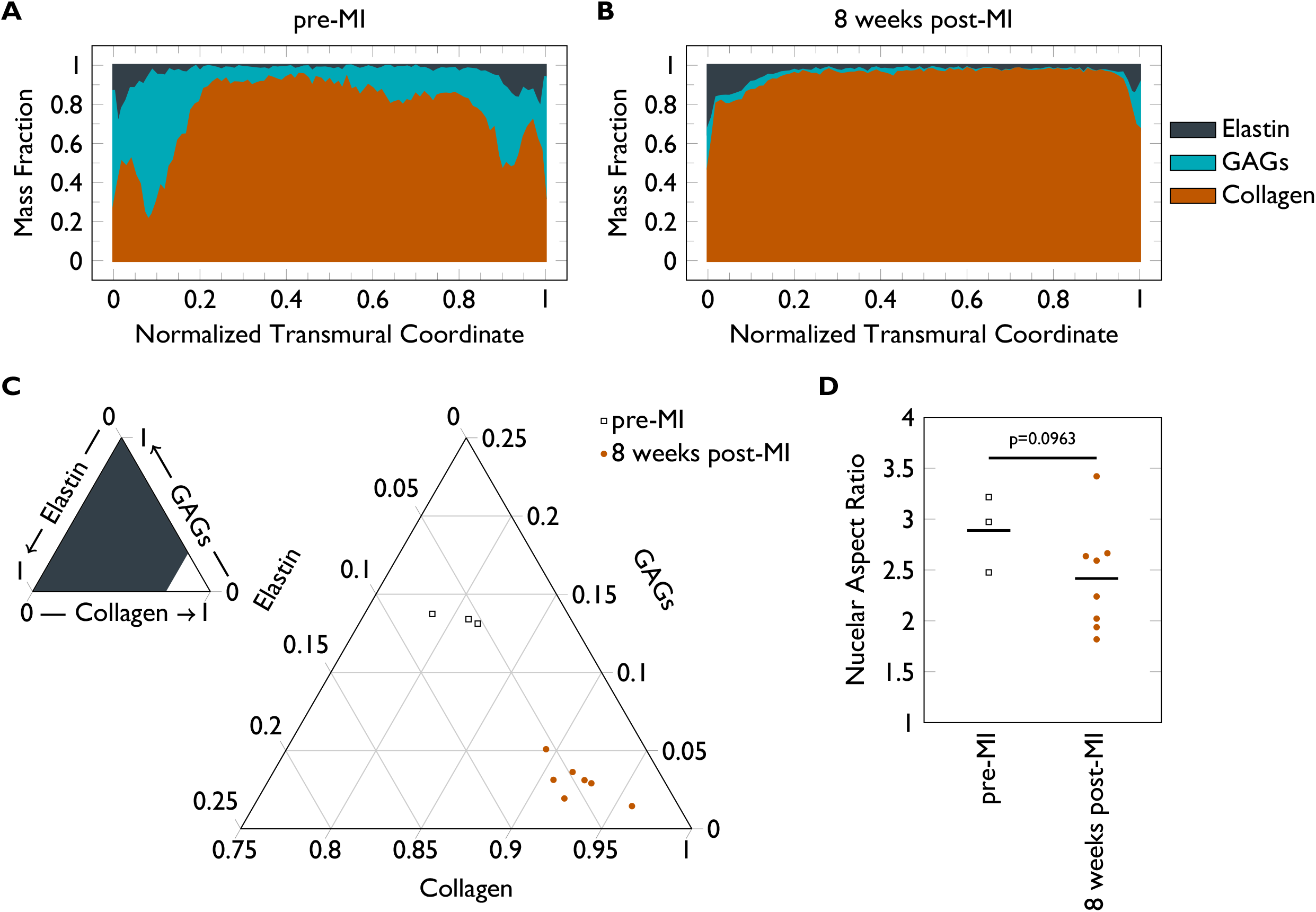
Leaflet compositional changes. Transmural distribution of ECM constituents (A) pre-MI versus (B) 8 weeks post-MI. (C) Overall composition change of pre-MI (n=3) versus 8 weeks post-MI (n=8). Groups compared with a Hotelling’s T^2^ test in the orthonormal coordinate system in 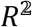 described in the Methods. (D) NAR changes pre-MI (n=3) versus 8 weeks post-MI (n=8). Groups compared with a Student’s t-test.

### 3. MVIC geometric changes

Such an irregular ECM structure and composition could affect MVIC geometry. In fact, Figure 4 alludes to a generally rounder MVIC at 8 weeks post-MI as compared to native tissue. Quantifying NAR from H&E stained slides confirms this observation with a decrease in observed NAR from 2.8883 ± 0.1025 to 2.4162 ± 0.1327 (p-value = 0.0963) at 8 weeks post-MI. This suggests that alterations at the tissue scale in both mechanical and material properties propagate down to mechanical microenvironment–induced MVIC geometric changes.

### 4. Gene expression

The observed disturbances in tissue mechanical properties, fiber architecture, and cell geometry in samples with MR at 8 weeks post-MI prompted us to investigate changes in the time course of gene expression in samples with and without MR post-MI. A second set of leaflets were processed for RNA-seq analysis, with samples available at pre-MI (without MR), 4 weeks post-MI without MR, 4 weeks post-MI with MR, 8 weeks post-MI without MR, and 8 weeks post-MI with MR. The data were first modeled as solely a function of time, resulting in 76 and 1 genes differentially expressed at 4 weeks and 8 weeks post-MI, respectively (Wald test, adjusted p-value < 0.1). Including the interaction between the presence of MR and the 4 weeks post-MI time point resulted in 25 genes better represented by the expanded model (LRT, adjusted p-value < 0.1); however, including the interaction between the presence of MR and the 8 weeks post-MI time point resulted in 0 genes better represented by the expanded model (LRT, adjusted p-value < 0.1). Therefore, the final model only included the interaction between the presence of MR at 4 weeks post-MI. The final model is formalized in Equation 2, where the counts *K_ij_* for gene i and sample j are modeled with a negative binomial distribution with mean *μ_ij_* and dispersion parameter α_*i*_. Mean μ_*ij*_ is composed of a term *q_ij_* proportional to the true concentration of reads and a sample-specific size factor *S_j_*. Finally, the coefficients in β^(*i*)^ give the log_2_ fold changes for gene i for each row of the model matrix ***X***, where entries in ***X*** are logical values in {0,1}. All of the genes with an adjusted p-value of ≤ 0.1 for 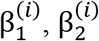, or 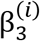 in the final model are presented in Figure 6. Since the interaction between the presence of MR and the 8 weeks post-MI time point did not contribute to the explanation of this data and relatively few genes displayed changes in gene expression at 8 weeks post-MI, most of the changes in gene expression due to MI have returned to baseline by 8 weeks.

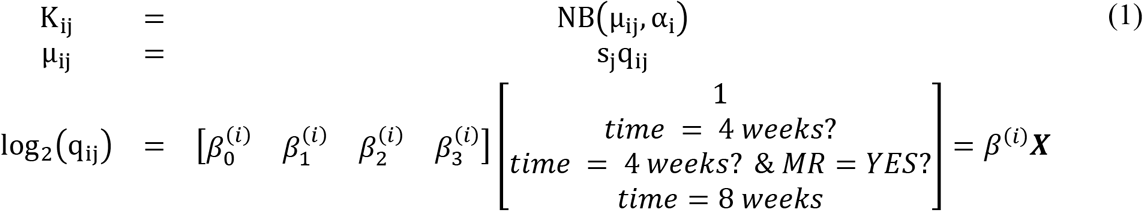

**Figure 6:**
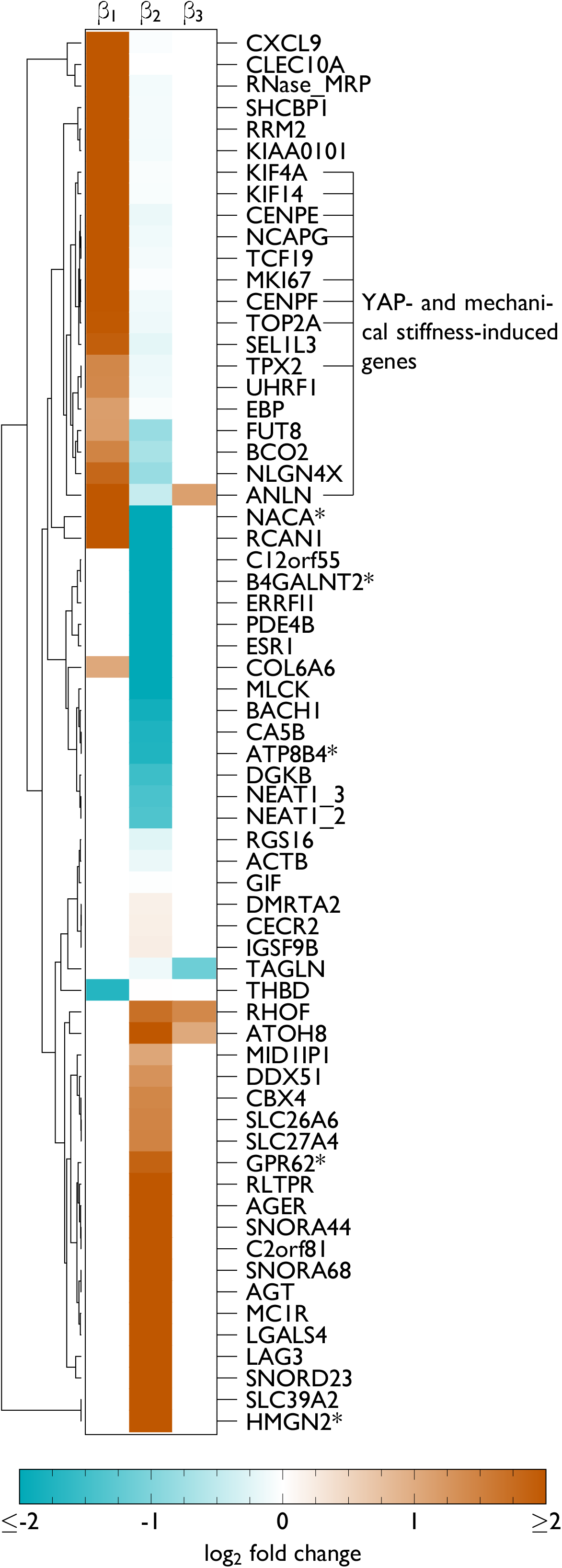
Heatmap of log_2_ fold changes for genes with a p-value of ≤ 0.1 (Wald test with Benjamini-Hochberg correction) for any parameter in the final model. Asterisks indicate gene names that were not present in the original annotation and instead found through BLAST searches on the Ovis aries genome.

Although large changes in expression for individual genes may significantly impact cellular processes, small changes in many genes within a single unifying biological process are often much more indicative of changes in cell phenotype.^50^ Furthermore, the list of significant genes often varies between research groups studying the same system, a problem exacerbated by small samples typical of exploratory RNA-seq experiments.^50^ Therefore, gene set enrichment analysis (GSEA)^50^ using gene sets pre-defined in the Reactome^51^ database was used to investigate significant collections of genes. Ranking all genes by *β*_1_, the differential expression due to 4 weeks post-MI alone, resulted in four pathways with significant alterations (adjusted p-value < 0.1); however, the core enrichment for all of these pathways consisted of a single gene, CXCL9.

Instead, ranking all genes by *β*_2_, the differential expression due to the presence of MR at 4 weeks post-MI, resulted in significant differences (adjusted p-value < 0.1) in 27 pathways (Supplemental Table 1), of which downregulation of “Extracellular Matrix Organization” and “ECM proteoglycans” were among the most significant differences at the level of gene expression. “Collagen chain trimerization”, “collagen biosynthesis and modifying enzymes”, and “collagen degradation” are also significantly downregulated, agreeing with the increased collagen mass fraction, but shorter fibers at the tissue level. Finally, “smooth muscle contraction” is significantly decreased in agreement with the decrease in NAR found through quantitative histology. Together, these gene-level results suggest that most changes in gene expression have returned to baseline by 8 weeks post-MI and changes in gene expression between samples with and without MR at 4 weeks post-MI are primarily tied to the changes observed in ECM synthesis and remodeling observed at the tissue-level.

## Discussion

The present study, along with the previously reported in vivo results^23^, characterize the multiscale nature of IMR as illustrated in Figure 7. At the organ level, MI causes LV wall thinning and dilation, tethering the MV leaflets. At the valve level, leaflet tethering induces a prominent plastic radial deformation that alters the leaflet biomechanical behavior. This plastic deformation is accompanied by decreased GAG and increased collagen content, with some evidence of shorter collagen fibers, at the tissue level. At the cell level, VICs appear slightly rounder and transcriptomics points to potential immune cell infiltration, in agreement with previous reports.^29^ Furthermore, others have observed endothelial mesenchymal transition (EndMT) in valve endothelial cells (VECs).^29^ Finally, at the gene level, YAP mechanotransduction is prominent at 4 weeks post-MI. Genes regulating ECM organization are downregulated in samples that have experienced IMR at 4 weeks post-MI compared to samples that did not experience IMR. Each scale is further discussed in the text that follows.

**Figure 7:**
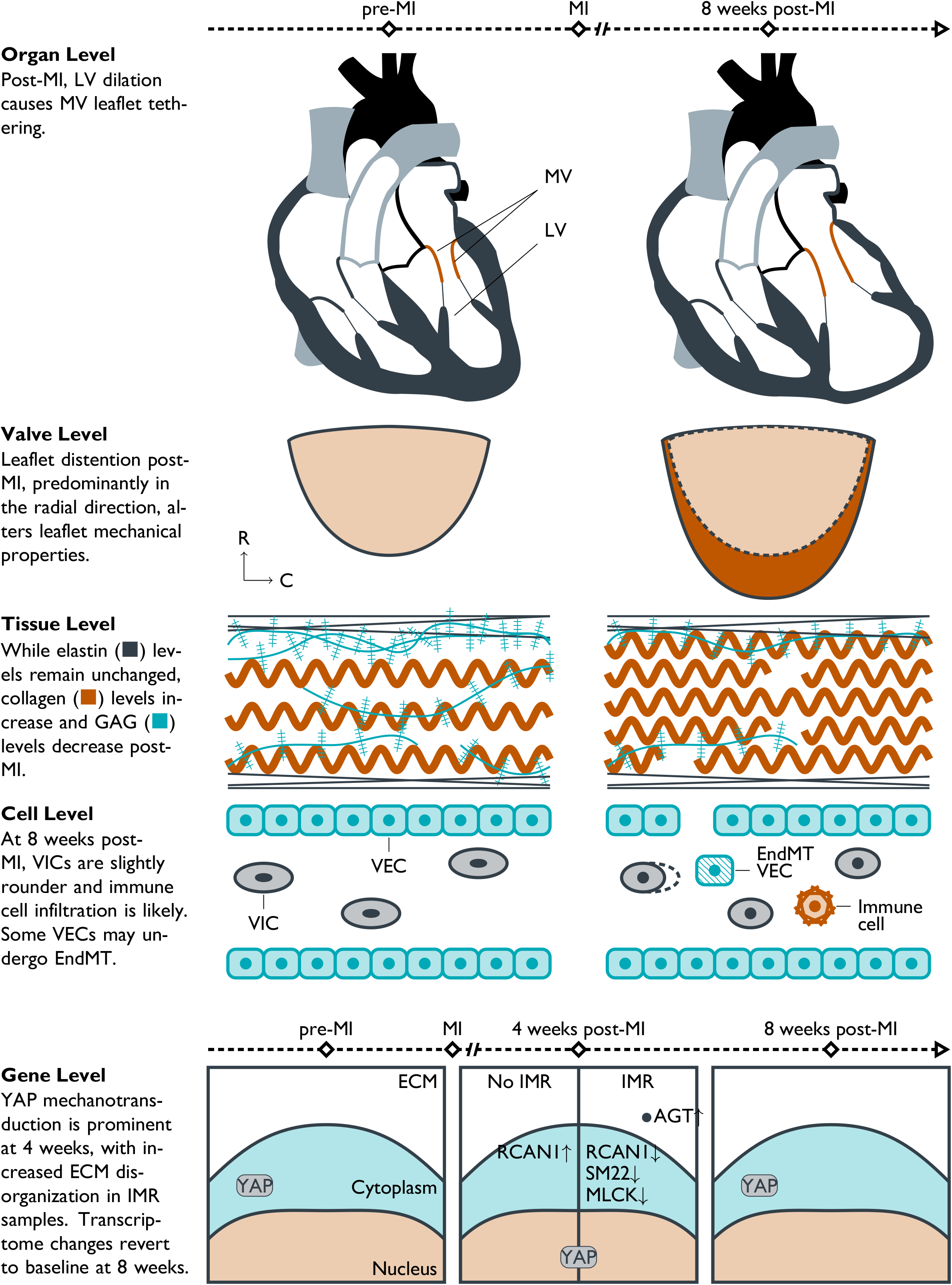
Illustration of the multiscale behavior of IMR.

### 1. Changes in mechanical behavior and composition are the result of permanent radial distention

The mechanics and composition of the MV were found to be substantially altered at 8 weeks post-MI. At 8 weeks post-MI, the radial peak strain significantly decreased, indicating a loss of mechanical anisotropy. These in vitro results are consistent with the in vivo conclusions that leaflet tethering has a larger effect on MV remodeling than annular dilation.^23^ Moreover, the changes in mechanical properties for the MV leaflet appear to be almost entirely attributable to *MI-induced permanent radial distention of the leaflet*, which causes the radial stiffness to increase substantially even in the absence of changes to the material stiffness of the underlying collagen fiber network. That is, radial distention resulting from persistent leaflet tethering causes the more radially-oriented fibers to straighten, and thus they are recruited at lower levels of strain with respect to the leaflet’s new unloaded configuration.^3,39,55^

Moreover, the observed reduction in MVIC nuclear aspect ratio (Figure 4 and Figure 5) are likely a direct result of the pronounced radial distension that occurred post MI (Figure 3). This is based on our previous work on quantitative relations between the MV leaflet stretch and local MVIC deformations^27,54^ where we quantitatively demonstrated MVIC deformation can be directly predicted from organ and tissue level deformations. Moreover, we identified MVIC NAR as a key indicator in leaflet tissue homeostatic regulation and, as such, can be used as a key metric in determining MVIC biosynthetic behavior. These studies indicated that MVIC responses have a delimited range of in vivo deformations that maintain a homeostatic response, suggesting that deviations from this range may lead to deleterious tissue remodeling and failure. However, future, longer-term investigations are necessary to determine whether active growth and/or remodeling play a substantial role in the mechanical adaptation of the MV to its post-MI ventricular environment beyond the initial 8-week period spanned by the present study.

### 2. Changes in gene expression reveal both MVIC and immune cell responses to IMR

RNA sequencing experiments provided unprecedented insight into the transcriptomic changes occurring in response to IMR. Angiotensin (AGT) is one of the most significantly and strongly upregulated genes in sheep experiencing IMR at 4 weeks (*β*_2_=2.38, adjusted p-value = 0.034), suggesting that the angiotensin-converting-enzyme inhibitors (ACEIs) and angiotensin receptor blockers (ARBs) commonly prescribed for coronary artery disease may have additional affects in the MV. In fact, patients tolerating maximal ACEI/ARB therapy exhibit reduced leaflet thickening post-MI.^56^ Regulator of calcineurin 1 (RCAN1) is elevated at 4 weeks post-MI (*β*_1_ = 2.09, adjusted p-value = 0.063), but decreased in the subset of sheep with MR at 4 weeks post-MI (*β*_2_ = −3.07, adjusted p-value = 0.0003). RCAN1 inhibits cardiac hypertrophy^57,58^ and is a direct transcriptional target of NFATc1 during valvulogenesis^59^. Overexpression of RCAN1 leads to reduced LV growth/remodeling at 4 weeks and reduced collagen deposition in a transgenic mouse model^58^ while silencing of RCAN1 increased pulmonary VEC migration in ovine post-natal valves^60^. Cyclosporine A and FK506 are pharmaceuticals that inhibit calcineurin and have profound effects on cardiac hypertrophy, but they were deemed inappropriate for treating cardiac hypertrophy due to nephrotoxicity.^61^ Myosin light chain kinase (MLCK) is downregulated at 4 weeks in sheep experiencing MR (*β*_2_ = −2.00, adjusted p-value = 0.040) and transgelin (TAGLN, SM22) is downregulated (*β*_3_= −1.14, adjusted p-value = 0.022) at 8 weeks post-MI, which agree with the decrease in NAR observed through histology. MKI67 is upregulated at 4 weeks post-MI (*β*_1_ = 2.06, adjusted p-value = 0.079), a marker previously used to indicate cell proliferation in IMR^29^. Finally, CXCL9 and CLEC10A are the most strongly upregulated gene at 4 weeks post-MI (CXCL9: *β*_1_ = 5.96, adjusted p-value = 0.003; CLEC10A: *β*_1_ = 5.01, adjusted p-value = 0. 079), indicating a possible attempt to recruit lymphocytes to the MV post-MI by dendritic cells^62^, similar to the role of CXCL9 in recruiting T cells to the MV in rheumatic disease^63^. The presence of LAG3 (*β*_2_ = 2.95, adjusted p-value = 0.005) in sheep exhibiting MR at 4 weeks post-MI also points to T cell infiltration^64^, possibly pointing to the same infiltrating cell population found previously^29^ through staining for CD45. Although there are limited studies investigating the immune response and inflammation cascade involved in the context of IMR, evidence of platelet activation has been observed in MR caused by MV prolapse^65,66^, which could also contribute to leukocyte recruitment and activation in IMR.

### 3. Altered mechanotransduction and ECM organization are major gene groups altered in response to IMR

In addition to roles of individual genes, grouping genes by promoter or function can provide further insight into the function of VICs in IMR. Many of the genes upregulated at 4 weeks post-MI were also found in a previous study comparing genes that are induced both by YAP in NIH 3T3 fibroblasts and mechanical stiffness in human lung fibroblasts^67^, indicating increased mechanical activation at 4 weeks post-MI and agreeing with the larger strains observed radially post-MI in vivo^23^. In the GSEA analysis, decreases in genes related to “Extracellular Matrix Organization” and “ECM proteoglycans” obtained by ranking all genes by *β*_2_ agree with protein/tissue level changes in the overall GAG component of the ECM and shorter collagen fibrils observed in quantitative histology and SEM, respectively. These results indicate remarkable consistency across all scales investigated.

### 4. Limitations of the present study

Although this study presented a detailed multiscale view into the mechanism of MV remodeling in IMR, limitations remain. First, IMR is an intrinsic disease of the LV, so the extent and location of the MI likely influence the observed MV remodeling. In the present study, we controlled for this effect by inducing posterobasal MIs of equal size in all animals. Although this type of MI has substantial clinical relevance, future studies should investigate the sensitivity of MV remodeling phenomena to other types of MI. Next, at the cellular level, there was no effort to separate VICs from valve endothelial cells (VECs) or infiltrating immune cells, and conclusions drawn about specific cell types are thus limited. For instance, a constant level of αSMA could either be because IMR did not affect VIC/VEC αSMA expression or an increase in VEC αSMA expression, as observed previously^29,31,56^, is balanced by a decrease in VIC αSMA expression. Therefore, the results were only attributed to specific cell types if the gene of interest has a very restricted cell type distribution (e.g., CLEC10A for CD1c+ dendritic cells^62^). The limitations of this study suggest future directions for characterizing MV remodeling in response to IMR.

### 5. Conclusions and future studies

The multiscale experiments conducted herein are crucial for informing advanced computational simulations of the MV post-MI.^26,68^ Mechanical testing of tissues post-MI revealed a critical change in the anisotropy observed in pre-MI valves, consistent with a substantial and permanent radial leaflet distention induced by leaflet tethering. The material composition was also altered, with more collagen, less GAGs, and similar elastin compositions. This data can better inform MV material models post-MI as one of the three critical pillars of MV in vivo simulation post-MI.^55,69,70^ Such computational models promise to suggest optimal device designs and patients suitable for one treatment over another. Moreover, connecting to the level of the transcriptome paves the way for multiscale models to extend down to signaling mechanisms with the goal of eventually evaluating both surgical and pharmaceutical interventions in one unified computational framework.

## Acknowledgments

None.

