## Supporting Information for "Mitral valve leaflet response to ischemic mitral regurgitation: From gene expression to tissue remodeling"

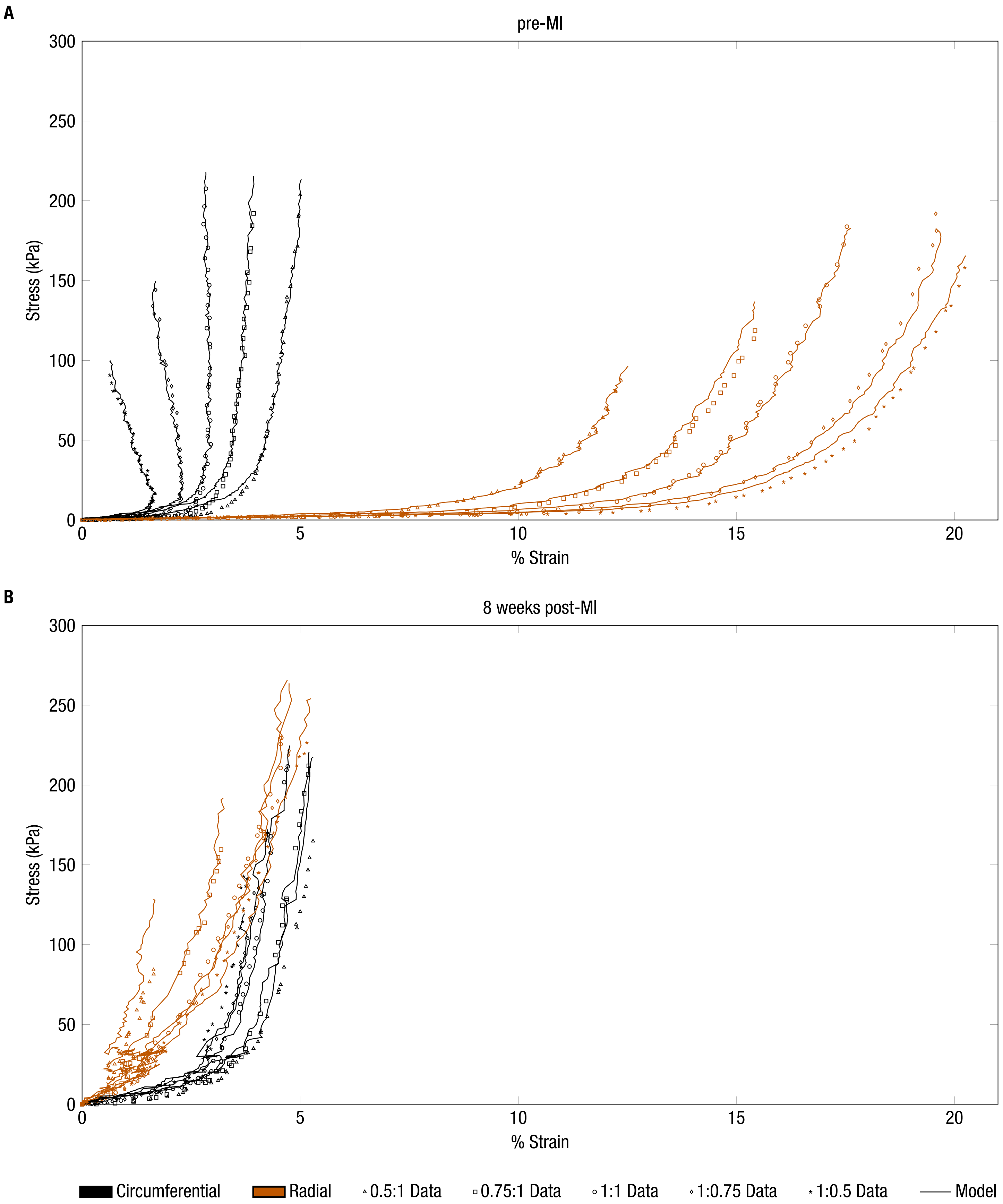


Supplemental Figure 1: Full biaxial mechanical testing results and model fits for representative specimens (A) pre-MI and (B) 8 weeks post-MI.

Supplemental Table 1: GSEA results using the Reactome database. The normalized enrichment score (NES) characterizes the up-/downregulation and the p-value is the adjusted p-value using the Benjamini-Hochberg criterion.

| ID | Description | NES | p-value |
|---|---|---|---|
| R-HSA-1474244 | Extracellular matrix organization | -1.729 | 0.037 |
| R-HSA-3000178 | ECM proteoglycans | -1.921 | 0.037 |
| R-HSA-111885 | Opioid Signalling | -1.867 | 0.037 |
| R-HSA-6809371 | Formation of the cornified envelope | 2.144 | 0.037 |
| R-HSA-5368287 | Mitochondrial translation | 1.936 | 0.037 |
| R-HSA-5389840 | Mitochondrial translation elongation | 1.948 | 0.037 |
| R-HSA-5368286 | Mitochondrial translation initiation | 1.942 | 0.037 |
| R-HSA-6805567 | Keratinization | 2.173 | 0.037 |
| R-HSA-381753 | Olfactory Signaling Pathway | -1.785 | 0.052 |
| R-HSA-5083625 | Defective GALNT3 causes familial hyperphosphatemic tumoral calcinosis (HFTC) | 1.995 | 0.053 |
| R-HSA-5419276 | Mitochondrial translation termination | 1.904 | 0.053 |
| R-HSA-3000171 | Non-integrin membrane-ECM interactions | -1.867 | 0.056 |
| R-HSA-8866910 | TFAP2 (AP-2) family regulates transcription of growth factors and their receptors | -1.872 | 0.056 |
| R-HSA-72312 | rRNA processing | 1.808 | 0.066 |
| R-HSA-375276 | Peptide ligand-binding receptors | 1.757 | 0.076 |
| R-HSA-6791226 | Major pathway of rRNA processing in the nucleolus and cytosol | 1.745 | 0.076 |
| R-HSA-5083636 | Defective GALNT12 causes colorectal cancer 1 (CRCS1) | 1.966 | 0.079 |
| R-HSA-8948216 | Collagen chain trimerization | -1.820 | 0.079 |
| R-HSA-8868773 | rRNA processing in the nucleus and cytosol | 1.785 | 0.079 |
| R-HSA-72766 | Translation | 1.655 | 0.079 |
| R-HSA-445355 | Smooth Muscle Contraction | -1.832 | 0.079 |
| R-HSA-72203 | Processing of Capped Intron-Containing Pre-mRNA | 1.680 | 0.085 |
| R-HSA-112314 | Neurotransmitter receptors and postsynaptic signal transmission | -1.634 | 0.086 |
| R-HSA-1650814 | Collagen biosynthesis and modifying enzymes | -1.725 | 0.086 |
| R-HSA-72172 | mRNA Splicing | 1.749 | 0.086 |
| R-HSA-73854 | RNA Polymerase I Promoter Clearance | 1.873 | 0.090 |
| R-HSA-1442490 | Collagen degradation | -1.745 | 0.096 |
